# Independently tunable subcortical oscillators enable prefrontal control of cortical coupling

**DOI:** 10.64898/2026.10.13.758762

**Authors:** Marjan Mozaffarilegha, Angeliki Kolaxi, Vassiliki Nikoletopoulou, Hiroshi T. Ito

## Abstract

Oscillatory coherence between brain regions varies flexibly with behavioral demands, yet the circuit mechanism underlying this control remains unclear. Theta rhythm, which coordinates prefrontal-hippocampal activity, is classically attributed to a pacemaker in the medial septum (MS), with the supramammillary nucleus (SuM) as its modulator. Here we show that SuM instead operates as an independently tunable theta oscillator. In rats performing a spatial alternation task, SuM theta frequency decreased before the choice point while MS theta frequency remained stable, reducing the SuM-MS frequency mismatch in both population spiking and local field potentials. The size of this mismatch predicted theta-rhythm coordination between the medial prefrontal cortex (mPFC) and hippocampal CA1 in both the alternation task and open-field foraging. Inhibiting direct mPFC input to SuM abolished the behavior-dependent modulation of SuM frequency and disrupted mPFC-CA1 coupling. These findings place a subcortical theta oscillator under top-down prefrontal control and identify the SuM–MS frequency mismatch as a control variable for interregional coordination.

## Introduction

The brain’s computational capacity is thought to rely on distributed and parallel processing, but this architecture creates a fundamental control problem. Anatomical connections between regions are largely stable, yet the functional interactions during perception, learning, and action often need to be reconfigured on timescales of hundreds of milliseconds to seconds. Oscillatory synchrony has long been proposed as one mechanism for such flexible coordination: by aligning cycles of excitation and inhibition across regions, synchrony can transiently modulate the effectiveness of communication between selected populations without changing anatomical connectivity (*1–8*). Across perceptual, attentional, and memory tasks, changes in interregional coherence accompany changes in effective coupling (*7, 9–12*), and experimental and modeling work indicates that the frequency relationship between oscillators determines whether and when they synchronize, and thus which populations can communicate (*10, 13–15*). However, while decades of studies have established correlations between oscillatory coupling and behavior, the key mechanistic question—how the coupling itself is controlled—remains largely unsolved. Without identifying such a control mechanism, it remains unclear whether oscillatory coupling is actively regulated or merely a passive byproduct of shared input.

In rodents, theta oscillations in the range of 6-10 Hz are prominent during navigation in the hippocampal-entorhinal circuit, organizing spike timing and population sequences supporting spatial coding and planning (*16–24*). Theta-rhythm coordination extends beyond this circuit. During spatial memory and decision-making tasks, medial prefrontal cortex (mPFC) neuron spikes are phase-locked to hippocampal theta rhythm, with enhanced theta coherence between mPFC and hippocampal CA1 local field potentials (LFPs), and the strength of this temporal coordination predicts successful performance (*25–28*). Circuit manipulations that disrupt hippocampal– prefrontal theta coordination also impair spatial working memory (*29*). Theta-rhythm coordination across mPFC, thalamic nucleus reuniens, and hippocampal CA1 supports the transfer of prospective trajectory information from mPFC to CA1 (*30, 31*), and similar coordination is also observed between the hippocampus and the striatum at trajectory choice points (*32, 33*). Theta coordination is thus recruited across regions depending on behavioral demands, raising the question of how an apparently unitary rhythm can achieve such selectivity.

Theta rhythm has been attributed to both cortical and subcortical systems, but the medial septum (MS) has classically been regarded as its main pacemaker: MS lesions or inactivation abolish hippocampal theta, and rhythmic MS activations entrain it in behaving animals (*16, 34–36*). In addition, the supramammillary nucleus (SuM) has been considered another subcortical component for the theta system. SuM neurons discharge rhythmically with hippocampal theta (*37, 38*), and manipulations of SuM alter hippocampal theta frequency (*38–41*). However, silencing SuM did not abolish hippocampal theta rhythm in behaving animals (*42, 43*), leading SuM to be regarded as a modulator of MS theta rather than an independent oscillator.

This view however assumes that the brain has a single theta rhythm shared across regions. Whether the brain instead sustains multiple independently tunable theta rhythms has received little attention, although several observations point to this possibility. First, theta has classically been divided into lower-frequency atropine-sensitive and higher-frequency atropine-insensitive ones (*16, 44*), indicating at least two mechanistically distinct theta processes. Second, hippocampal theta frequency is dissociated during decision-making between dorsal and ventral regions (*45*). Third, silencing SuM spared local theta power in mPFC and CA1 yet disrupted mPFC–CA1 theta coordination at the choice point (*30*), which is not predicted by an account in which SuM merely modulates MS theta.

Multiple independent oscillators provide another level of control over interregional synchrony. Beyond merely being engaged or disengaged together, they can operate independently by changing their relative frequencies, which drives a continuous drift in their relative phases, so that downstream populations are entrained to each oscillator to varying degrees. Experimental and theoretical studies indicate that oscillatory frequency gates interregional communication (*7, 10, 13–15*). Whether the brain actively regulates the frequency relationship between multiple oscillators to control large-scale network coordination, however, remains unknown. Addressing this requires simultaneous recordings from SuM and MS, to determine whether their theta rhythms vary independently with behavioral demand, and manipulation of their afferents, to test whether this relationship is under upstream control.

Here we combined simultaneous recordings from SuM, MS, mPFC, and CA1 with pathway-specific optogenetic silencing of prefrontal afferents to SuM in freely behaving rats. We found that MS theta frequency was largely stable, whereas SuM theta frequency varied with the demands of trajectory decisions. This modulation was evident in both population spiking and LFPs and was independent of the animal’s speed or acceleration, pointing to control by cognitive demand rather than locomotion. This modulation reduced the frequency offset between the two oscillators and was accompanied by increased theta-rhythm coordination between their downstream targets, mPFC and CA1. We identified a direct cortical projection from the mPFC to SuM, and optogenetic silencing of its terminals abolished the modulation of SuM theta frequency and mPFC-CA1 coordination during trajectory decisions. These findings show that prefrontal input actively regulates the frequency relationship between independently tunable subcortical oscillators, providing a mechanism for flexibly organizing communication across distributed circuits.

### SuM theta is retuned to match MS theta before trajectory decisions

To determine how subcortical theta oscillations vary with trajectory-decision demands, we simultaneously recorded LFPs from SuM and MS while rats performed a spatial alternation task on a modified T-maze (11 recording sessions from 4 rats; Fig. 1A, C). In this task, rats ran along the stem part of the maze and chose a left or right turn at the T-junction to reach one of two goal wells. After consuming a water reward, they returned to the start well along a side arm (the nonstem segment). The rewarded goal well alternated between left and right on successive trials. The maze was designed so that the stem and nonstem segments followed geometrically identical paths.

**Fig. 1.**
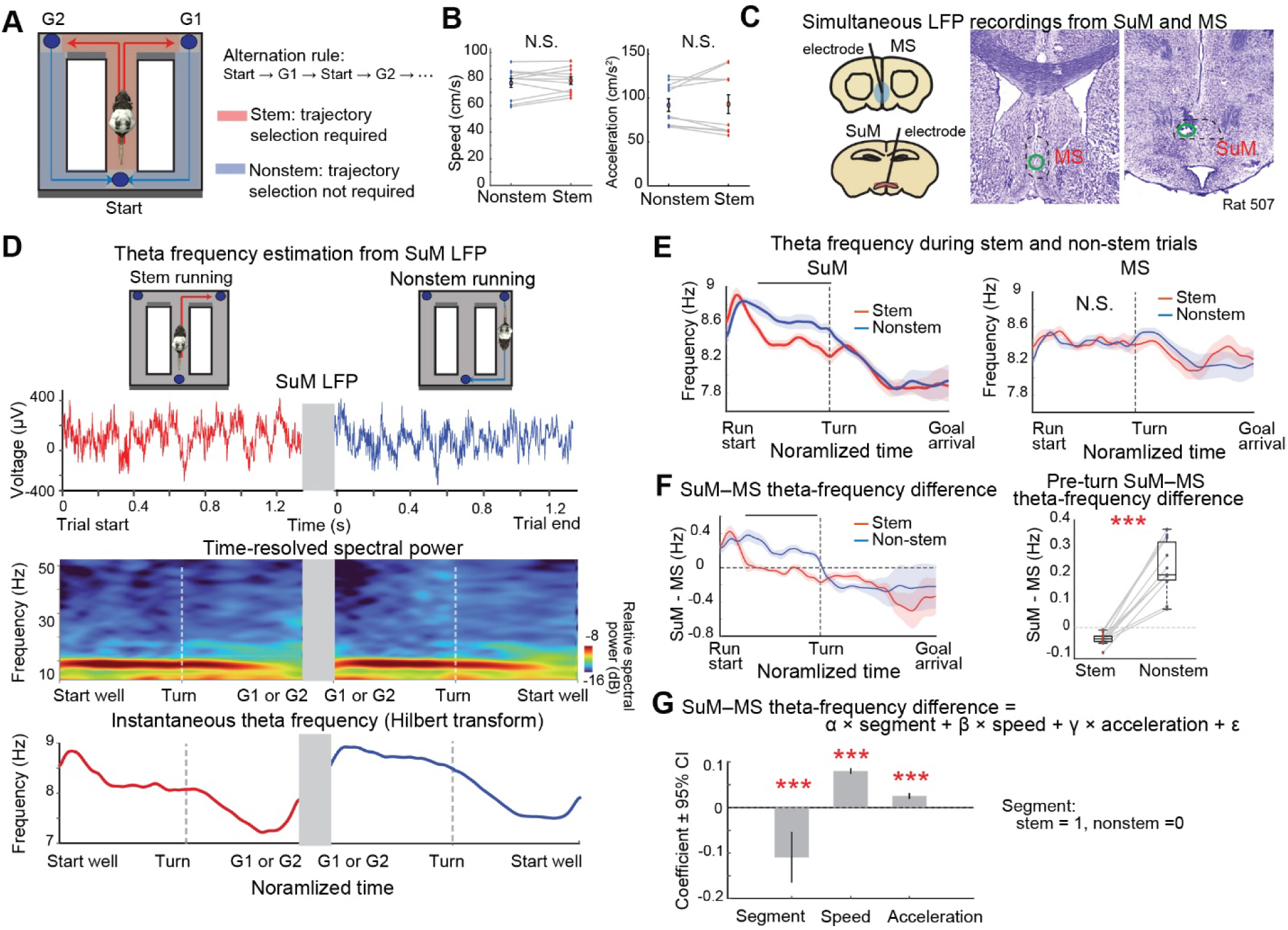
Modulation of SuM theta frequency and its offset from MS frequency before trajectory decisions. **(A)** Schematic of the modified T-maze alternation task. Rats alternated between Goal 1 (G1) and Goal 2 (G2) from the start well. Stem traversals preceded the trajectory choice, whereas nonstem traversals required no choice. **(B)** Running speed and acceleration across the pre- and post-turn phases of stem and nonstem traversals. Gray lines connect paired values from individual sessions. Speed and acceleration did not differ between stem and nonstem traversals (speed: p = 0.291, acceleration: p = 0.777, linear mixed-effects models; 11 sessions, 4 rats). **(C)** Left, schematic showing electrode placement in SuM and MS for simultaneous LFP recordings. Right, representative histological sections showing electrode tracks in MS and SuM. **(D)** LFP from SuM in a representative session, with its time-resolved power spectrum and instantaneous frequency. Power was estimated from the raw LFP and shows a prominent theta peak. Instantaneous frequency was obtained from the temporal derivative of the unwrapped Hilbert phase of the theta-band-filtered signal. **(E)** SuM and MS theta frequency across the two segments of normalized time, from start to turn and from turn to goal arrival. Lines and shading, session mean ± s.e.m. (11 sessions, 4 rats). Before the turn, SuM theta frequency was lower during stem than nonstem traversals (difference −0.25 Hz, p < 0.001), whereas MS frequency did not differ (+0.02 Hz, p = 0.187). The black bar marks bins with FDR-adjusted q < 0.01 in a linear mixed-effects model. After the turn, neither SuM (−0.01 Hz, p = 0.546) nor MS (−0.03 Hz, p = 0.328) differed between traversal types. Linear mixed-effects models with animal identity as a random effect throughout. **(F)** SuM–MS theta-frequency difference (Δf = fSuM − fMS) across normalized time before and after the turn. Before the turn, Δf was smaller during stem than nonstem traversals; the black bar marks bins with FDR-adjusted q < 0.01 in a linear mixed-effects model. Right, paired session averages: Δf was −0.04 ± 0.01 Hz during stem and +0.22 ± 0.03 Hz during nonstem traversals (p < 0.001). After the turn, Δf did not differ between traversal types (p = 0.557). **(G)** Generalized linear model of the SuM–MS frequency difference across pre- and post-turn phases. Bars show fixed-effect coefficients and 95% confidence intervals for segment (stem/nonstem), running speed, and acceleration. Segment remained a significant predictor (−0.109 Hz, p < 0.001) when running speed (0.080 Hz per cm/s, p < 0.001) and acceleration (0.025 Hz per cm/s^2^, p < 0.001) were included in the same model.

Because theta frequency covaries with running speed and acceleration (*46, 47*), we first compared locomotion between the stem and nonstem segments. Running speed and acceleration did not differ significantly between stem and nonstem traversals (speed: stem 78.89 ± 2.80 cm/s, nonstem 77.03 ± 3.33 cm/s, p = 0.291; in a linear mixed-effects model with animal identity as a random variable, 4 rats; acceleration: stem 92.98 ± 10.67 cm/s², nonstem 91.90 ± 7.20 cm/s², p = 0.777; Fig. 1B).

We then examined the LFPs recorded from SuM and MS while the rat performed this task (Fig. 1D). Time-resolved spectral power analysis of LFPs revealed a prominent peak near the theta band in both regions. To estimate an instantaneous theta frequency, we bandpass-filtered LFPs at 6–12 Hz, extracted phases by Hilbert transform, and computed frequency as the rate of phase advance.

Theta frequencies in SuM and MS LFPs were modulated differently during the task (Fig. 1E). Before the turn, SuM theta frequency was lower than during nonstem traversals (frequency: stem 8.38 ± 0.05 Hz, nonstem 8.63 ± 0.06 Hz; p < 0.001, linear mixed-effects model). MS theta frequency did not differ significantly between the same segments (stem 8.42 ± 0.05 Hz, nonstem 8.41 ± 0.05 Hz; p = 0.187). Neither region showed a significant difference after the turn (SuM: p = 0.546; MS: p = 0.328), indicating that theta-frequency modulation was specific to SuM and to the period preceding the trajectory choice.

This SuM-specific modulation of theta frequency altered the frequency offset between the two oscillators. Before the turn, the SuM-MS frequency offset was significantly smaller during stem than during nonstem traversals, with the two oscillators nearly matched in frequency (SuM-MS frequency difference: stem −0.04 ± 0.01 Hz, nonstem +0.22 ± 0.03 Hz; p < 0.001; Fig. 1F). No such difference was observed after the turn (p = 0.557).

We next used a generalized linear model (GLM) to test whether segment (stem vs. nonstem), running speed, and acceleration independently accounted for variation in the SuM–MS frequency offset across pre- and post-turn phases (Fig. 1G). This analysis revealed that the segment remained a significant predictor after accounting for speed and acceleration, both of which also contributed (stem/nonstem segment: α = −0.109, p < 0.001; speed: β = 0.080, p < 0.001; acceleration: γ = 0.025, p < 0.001). We further confirmed this result by subsampling trials so that mean running speed was matched between stem and nonstem segments; the SuM–MS frequency offset still differed significantly between segments in this subset (p < 0.001, linear mixed-effects model). These results together confirmed that SuM-MS offset was modulated by the task demand difference between stem and nonstem beyond the effect of locomotion.

Because the change in frequency difference was small in magnitude (0.26 Hz between stem and nonstem), we asked whether it was reliable relative to trial-to-trial variability. We randomly split stem and nonstem trials into two halves each. We then compared differences in frequency offset within segments (between halves of the same segment) with differences across segments. Across-segment differences were significantly larger than within-segment differences (median within-state difference, 0.07 Hz; median between-state difference, 0.26 Hz; p = 0.001). The segment effect therefore exceeded the variability expected from trial sampling alone.

To estimate the functional impact of this small change in frequency offset, we simulated two oscillators with varying frequency offsets and computed their phase coherence. The simulation window was 1.62 s, matching the mean pre-turn traversal time of stem and nonstem segments (stem, 1.62 ± 0.01 s; nonstem, 1.62 ± 0.01 s). Over this window, the observed change in offset (0.265 Hz, from nonstem [+0.222 Hz] to stem [−0.043 Hz]) increased phase coherence by 0.19. This is comparable to or larger than behavior-dependent changes in oscillatory coherence reported previously (*25, 28*).

Finally, to test whether these results depended on the method used to estimate instantaneous frequency, we repeated the analysis using the synchrosqueezed wavelet transform (*48*). Unlike the Hilbert-based approach, which requires narrow band-pass filtering, synchrosqueezing reassigns wavelet coefficients to sharpen the time–frequency representation, which gives high frequency resolution from short data segments without prior band-pass filtering. This method yielded the same conclusion: the SuM–MS frequency offset differed significantly between stem and nonstem segments (stem 0.00 ± 0.01 Hz; nonstem 0.25 ± 0.03 Hz; p < 0.001, linear mixed-effects model). SuM theta frequency also differed significantly between segments (stem 8.46 ± 0.053 Hz; nonstem, 8.70 ± 0.07 Hz; p < 0.001), whereas MS theta frequency did not (stem 8.45 ± 0.05 Hz; nonstem, 8.45 ± 0.05 Hz; p = 0.824).

Together, these results show that SuM theta frequency was modulated independently of MS before trajectory decisions, reducing the frequency mismatch between the two oscillators and bringing them into a more synchronized state. Because this effect was confined to the pre-turn phase, we restricted subsequent analyses to this phase of stem and nonstem traversals.

### SuM neurons reduce the frequency of theta-rhythmic spiking before trajectory decisions

To determine whether the segment-dependent modulation of SuM theta frequency was expressed in local spiking, we examined single-neuron rhythmicity in simultaneously recorded SuM and MS populations during the same stem and nonstem traversals (1,338 SuM neurons and 1,080 MS neurons, 11 recording sessions from 4 rats; Fig. 2A,B).

**Fig. 2.**
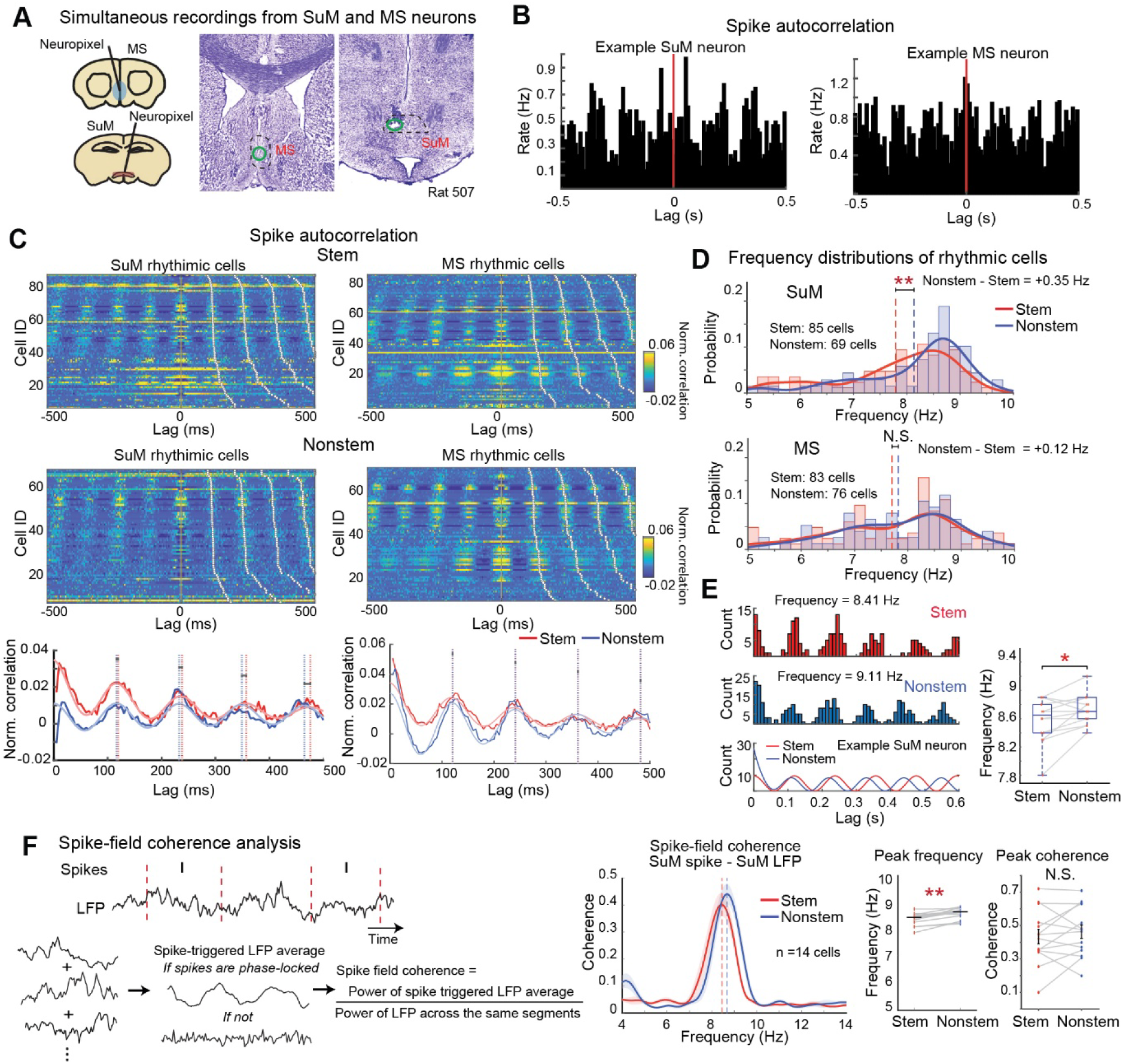
SuM neurons shift their rhythmic firing frequency with trajectory-decision demand. **(A)** Left, schematic of simultaneous Neuropixels recordings from SuM and MS. Right, histological sections showing probe tracks in MS and SuM. **(B)** Representative spike autocorrelograms from one SuM and one MS neuron, showing theta-timescale rhythmicity. **(C)** Autocorrelograms of theta-rhythmic SuM and MS neurons during stem and nonstem traversals. Heat maps show each neuron’s normalized spike autocorrelation, ordered by estimated rhythmic frequency within each condition (85 SuM stem and 69 SuM nonstem neurons; 83 MS stem and 76 MS nonstem neurons, 11 sessions, 4 rats, identified by a shuffle-based criterion applied to each condition separately; see Methods). Bottom, population-averaged autocorrelation profiles for stem (red) and nonstem (blue) traversals. In SuM, successive fitted peaks diverge progressively between conditions, whereas MS peak timing remains aligned. **(D)** Distribution of autocorrelogram-derived rhythmic frequencies across theta-rhythmic neurons. In SuM, frequencies were lower during stem (7.79 ± 0.12 Hz, n = 85 neurons) than during nonstem traversals (8.13 ± 0.12 Hz, n = 69; p = 0.041, two-sample Kolmogorov–Smirnov test). In MS, the distributions did not differ (stem 7.70 ± 0.13 Hz, n = 83; nonstem 7.82 ± 0.12 Hz, n = 76; p = 0.918). Values are mean ± s.e.m.; smooth curves are kernel density estimates. **(E)** SuM neurons classified as rhythmic in both stem and nonstem traversals (n = 14). These neurons shifted their rhythmic firing from 8.58 ± 0.08 Hz during stem to 8.73 ± 0.06 Hz during nonstem traversals (p = 0.014, linear mixed-effects model). Gray lines connect measurements from the same neuron. **(F)** Spike-field coherence between SuM LFP and SuM neurons rhythmic in both stem and nonstem traversals (n = 14 neurons). Left, schematic of the analysis: the spectral power of the spike-triggered average was normalized by the power of the same LFP segments. Middle, spike– field coherence for stem and nonstem traversals, showing a shift in peak frequency with comparable coherence magnitude. Right, frequency of maximal coherence differed between conditions (stem: 8.45 ± 0.07 Hz; nonstem: 8.68 ± 0.06 Hz; p < 0.001, linear mixed-effects model), whereas peak coherence did not (stem: 0.43 ± 0.04; nonstem: 0.46 ± 0.04; p = 0.350, linear mixed-effects model). Gray lines connect measurements from the same neuron. Coherence was computed from simultaneously recorded spikes and local SuM LFP with trial numbers balanced between conditions; peak frequency was estimated within 5–10 Hz.

Using a maximum-likelihood model of spike-train rhythmicity (*49*) (see Methods), we identified 10.46% of SuM neurons and 12.41% of MS neurons as significantly theta-rhythmic. Rhythmicity was often segment-specific, with many neurons rhythmic in only one segment (rhythmic during stem/nonstem: SuM, n = 85/69; MS, n = 83/76).

Color-coded spike autocorrelograms of rhythmic neurons showed that theta-band intervals in SuM were overall longer during stem than nonstem traversals, corresponding to a lower rhythmic frequency and consistent with the LFP results (Fig. 2C). MS neurons showed no such difference. Quantification of single-neuron rhythm frequency confirmed this: among segment-specific rhythmic neurons, the distribution of SuM theta frequencies was shifted toward lower frequencies during stem traversals (stem 7.79 ± 0.12 Hz, nonstem 8.13 ± 0.12 Hz; p = 0.041, Kolmogorov– Smirnov test; Fig. 2D). The MS distribution did not differ between segments (stem 7.70 ± 0.13 Hz, nonstem 7.82 ± 0.12 Hz; p = 0.918). The segment-dependent frequency modulation observed in the SuM LFP was thus accompanied by a corresponding shift in the population rhythmicity across SuM neurons, whereas MS neurons showed no comparable population-wide redistribution.

Because some neurons were rhythmic in only one segment, the stem and nonstem populations in Fig. 2D did not consist of the same cells. The frequency shift in SuM could therefore reflect different subsets of theta-rhythmic neurons in each segment, or a change in rhythmic frequency within the same neurons. To test the latter, we analyzed SuM neurons that were significantly rhythmic during both stem and nonstem traversals (n = 14 neurons). These neurons had a lower rhythmic frequency during stem than nonstem traversals (stem 8.58 ± 0.08 Hz, nonstem 8.73 ± 0.06 Hz; p = 0.014, linear mixed-effects model; Fig. 2E), suggesting that behavioral demands can change the intrinsic rhythmicity of individual neurons.

Finally, we asked whether the coupling between SuM spiking and local LFP reflected the same frequency shift between segments. To test this, we computed spike-field coherence from spike-triggered LFP averages (*50, 51*) (see Methods). SuM neurons that were rhythmic in both segments showed significant theta-band spike-field coherence, and coherence magnitude did not differ between stem and nonstem traversals (stem, 0.43 ± 0.04; nonstem, 0.46 ± 0.04; p = 0.350, linear mixed-effects model; Fig. 2F). Spike-theta coupling was therefore maintained across segments. The peak frequency of spike-field coherence, however, was lower during stem than nonstem traversals (stem, 8.45 ± 0.07 Hz; nonstem, 8.68 ± 0.06 Hz; p < 0.001; Fig. 2F). Thus, SuM neurons remained coupled to local theta while the frequency of this coupling shifted with behavioral demand.

Together, these results show that SuM and MS spiking mirrored the frequency relationships observed in the LFP. MS neurons maintained their rhythmic frequency across segments, whereas the rhythmic frequency of SuM neurons decreased during stem relative to nonstem traversals. Among neurons rhythmic in both segments, this shift occurred without a change in spike-field coherence magnitude, indicating retuning rather than altered theta coupling. Theta frequency modulation before trajectory choice was therefore expressed at the level of SuM neurons.

### Theta retuning is expressed across SuM, but not MS, neuronal populations

Although a subset of SuM and MS neurons fired theta-rhythmically, a substantial fraction did not reach significance by autocorrelation-based criteria. This could reflect either genuinely theta-independent firing or too few spikes for reliable frequency estimation. Autocorrelation-based methods infer frequency from the periodicity of spike-time autocorrelograms, which requires sufficient spike pairs at cycle-length lags. A neuron that fires sparsely or skips cycles therefore yields a weak or ambiguous autocorrelogram peak, even if its spikes are phase-locked to theta; such neurons may nonetheless contribute to theta generation. We therefore estimated rhythmic frequency at the population level, without classifying individual neurons. For each neuron, we computed spike phases relative to reference sinusoids from 6 to 10 Hz (0.1-Hz steps) and calculated the phase differences between pairs of spikes (Fig. 3A; Methods). A neuron that fires rhythmically at a given frequency emits spikes at a consistent phase of the reference sinusoid at that frequency, so its pairwise phase differences cluster around 0°, whether or not it fires in every cycle. We pooled these differences across simultaneously recorded neurons and used their concentration as a measure of population tuning to each frequency. This measure requires no LFP reference and is not biased by spike count, allowing segments with different numbers of spikes to be compared.

**Fig. 3.**
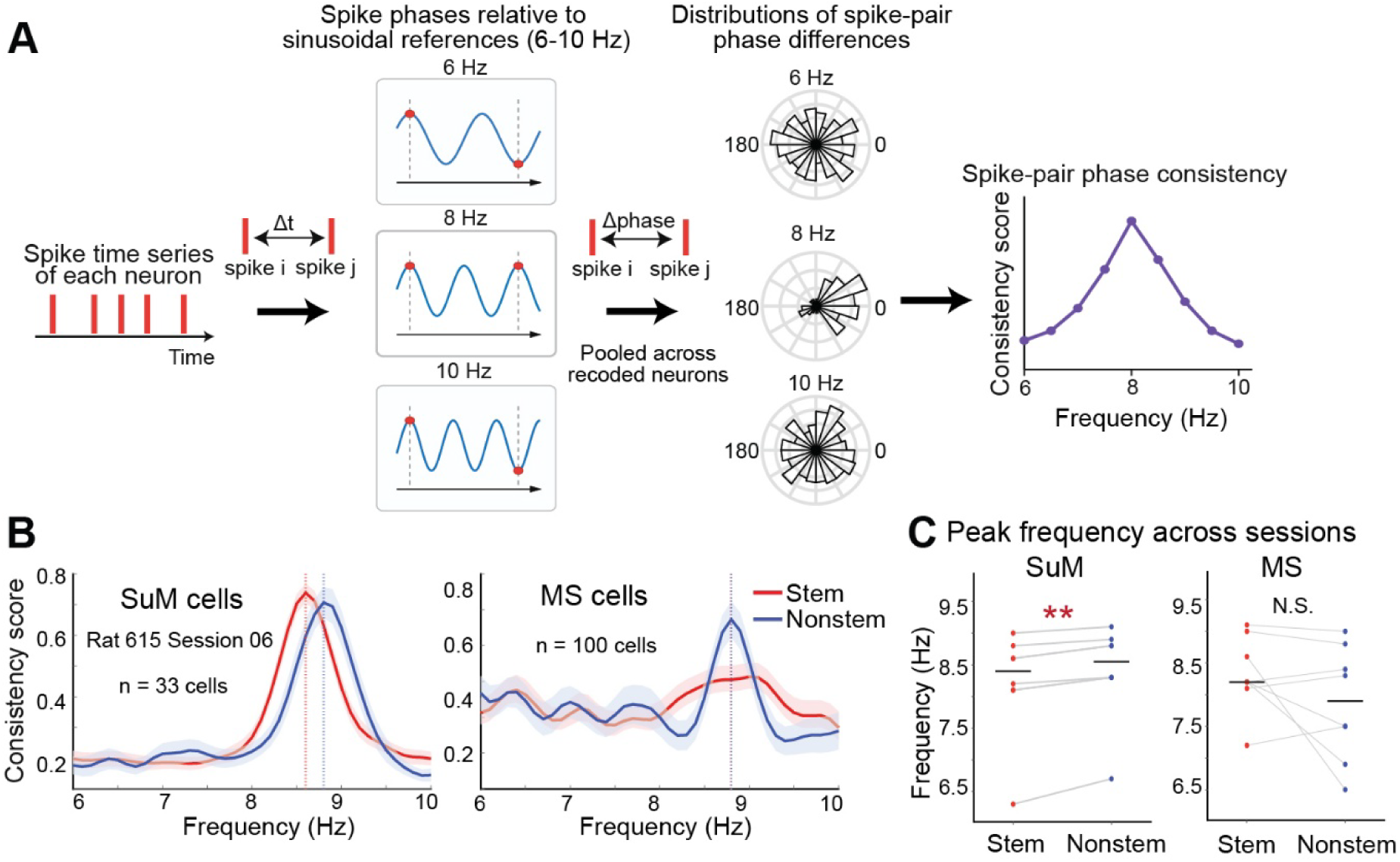
Trajectory-decision-dependent shift in rhythmic firing frequency across the SuM population. **(A)** Strategy for constructing population-level frequency-tuning profiles. For each neuron, every spike-pair interval was converted to a phase difference relative to reference sinusoids spanning 6–10 Hz in 0.1-Hz steps. Within each trial, phase-consistency values were pooled across simultaneously recorded neurons, weighted by the number of valid spike pairs. The resulting profile was min–max normalized for each trial and averaged across trials to obtain the population frequency profile. At a frequency matching the population rhythmicity, spike pairs fall at similar phases, and the phase differences cluster near 0°, without the need for neurons to fire on every cycle. **(B)** Representative population frequency profiles from a single recording session. Curves show the mean ± s.e.m. of normalized phase-consistency profiles across trials (33 SuM and 100 MS neurons) during stem (red) and nonstem (blue) traversals. The SuM population peak shifted from 8.6 Hz during stem to 8.8 Hz during nonstem traversals, whereas the MS peak was 8.8 Hz in both conditions. Dotted lines, peak frequencies. **(C)** Population rhythmic frequency during stem and nonstem traversals, shown separately for SuM and MS (8 sessions, 5 rats). Gray lines connect estimates from the same session. In SuM, population rhythmic frequency was lower during stem than nonstem traversals (difference −0.188 Hz, p < 0.001), whereas MS frequency did not differ between traversal types (+0.463 Hz, p = 0.148). Differences are stem minus nonstem; linear mixed-effects models with animal identity as a random effect.

In a representative session with simultaneously recorded SuM and MS neurons (Fig. 3B), the SuM population peak was lower during stem (8.6 Hz) than during nonstem (8.8 Hz) traversals, whereas the MS peak was 8.8 Hz in both segments, recapitulating the segment-dependent shift in single-neuron rhythmicity (Fig. 2). Across all sessions (8 sessions from 5 rats; Fig. 3C), SuM population rhythmic frequency was lower during stem traversals (stem 8.21 ± 0.30 Hz, nonstem 8.40 ± 0.27 Hz; p < 0.001, linear mixed-effects model). MS population frequency did not differ between segments (stem 8.33 ± 0.21 Hz, nonstem 7.86 ± 0.32 Hz; p = 0.148). The full width at half maximum (FWHM) of the SuM population peak did not differ between segments (stem 0.70 ± 0.02 Hz; nonstem, 0.77 ± 0.03 Hz; p = 0.062, linear mixed-effects model). Similarly, MS peak width showed no significant difference between segments (stem, 0.94 ± 0.08 Hz; nonstem, 1.20 ± 0.35 Hz; p = 0.453), and the distribution stayed predominantly unimodal in both (Hartigan’s dip tests of trial-wise peak frequencies pooled across sessions; SuM: stem, p = 0.701; nonstem, p = 0.949; MS: stem, p = 0.941; nonstem, p = 1.000). This indicates that the SuM population shifted as a whole toward lower frequency, rather than rhythmic neurons being redistributed between discrete frequency bands.

Together, these results extend the single-neuron findings to the population level: SuM population spiking was retuned between stem and nonstem traversals, whereas MS population spiking was not. SuM-specific retuning of population spiking thus reshaped the theta relationship between the two oscillators according to behavioral demand.

### mPFC input to SuM is required for decision-dependent retuning of SuM theta

Our results so far indicate that retuning of SuM neurons underlies the decision-dependent modulation of the SuM–MS frequency offset. How SuM neurons are informed of upcoming trajectory decisions, however, is unclear. A previous tracing study reported that retrograde tracer injected into SuM labeled neurons in mPFC, particularly in the infralimbic cortex, whereas anterograde tracer injected into the infralimbic cortex labeled only sparse fibers in SuM (*52*). The mPFC is a candidate source of this signal as it represents upcoming trajectories and conveys this information to CA1 through the nucleus reuniens (*30, 31*). We therefore examined whether mPFC axons form putative synaptic contacts in SuM, using anterograde tracing combined with immunostaining for synaptic markers and confocal microscopy.

We injected AAV-hSyn1-mApple into mPFC, targeting its full dorsoventral extent. mApple-labeled fibers were present in SuM, as well as in known mPFC targets, including the mediodorsal thalamic nucleus and nucleus reuniens (Fig. 4A). At higher magnification, mApple-labeled mPFC axons in SuM contained Bassoon-positive presynaptic puncta apposed to PSD-95-positive postsynaptic puncta, indicating that mPFC axons form putative excitatory synaptic contacts within SuM.

**Fig. 4.**
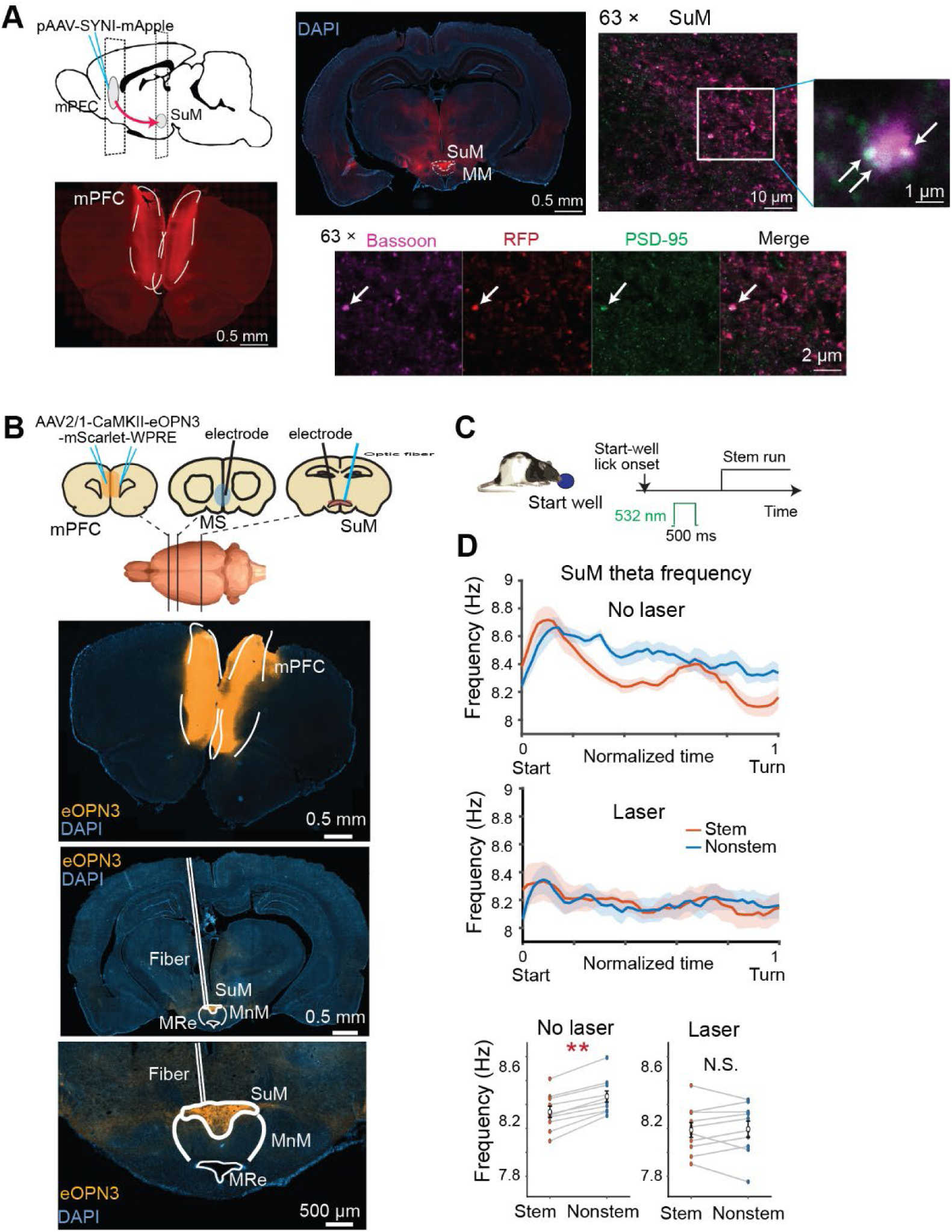
Silencing mPFC input to SuM impairs the trajectory-decision-dependent modulation of SuM theta frequency. **(A)** Anterograde tracing of mPFC projections to SuM. Top left, tracing strategy: a fluorescent reporter was expressed in mPFC neurons to label their axons in SuM. Bottom left, representative section showing reporter expression in mPFC. Middle and right, low- and high-magnification images of SuM showing labeled mPFC axons immunostained for Bassoon, a presynaptic active-zone marker, and PSD-95, a postsynaptic density marker. Insets, Bassoon-positive puncta within labeled mPFC axons apposed to PSD-95-positive puncta (arrows), consistent with putative synaptic sites between mPFC terminals and SuM neurons (9 sections from 3 rats). **(B)** Optogenetic silencing of mPFC terminals in SuM with simultaneous LFP recordings from SuM and MS. Top, strategy. Bottom, histological verification of, from top to bottom, eOPN3 expression in mPFC, optic fiber placement above SuM and labeled mPFC axon terminals in SuM. **(C)** Laser application protocol for eOPN3 activation. A single 500-ms pulse of 532-nm light was delivered to SuM between the onset of licking at the start well (end of the preceding trial) and the onset of the next trial. **(D)** SuM theta frequency in the pre-turn phase under laser-off and laser-on conditions. Top, SuM frequency over normalized time from start to turn; lines and shading, mean ± s.e.m. across stem and nonstem traversals. Bottom, session averages. With the laser off, SuM theta frequency was lower during stem than nonstem traversals (stem 8.34 ± 0.05 Hz, nonstem 8.47 ± 0.05 Hz; p < 0.001). During silencing of mPFC terminals, the two did not differ (stem 8.19 ± 0.06 Hz, nonstem 8.20 ± 0.07 Hz; p = 1.00). Linear mixed-effects model; 10 sessions, 4 rats.

To test whether mPFC input provides decision-dependent control of SuM, we expressed the inhibitory opsin eOPN3 (*53*) in mPFC neurons by injecting AAV-CaMKIIα-eOPN3 and implanted an optic fiber 500 µm above SuM. In the same animals, we also implanted LFP electrodes in SuM and MS. Because eOPN3 suppresses synaptic release with slow offset kinetics on the order of minutes (*53*), we delivered a single 500-ms pulse of 532-nm light before the start of each trial, which also avoided photoelectric artifacts in the LFP recordings. Each trial comprised a stem traversal followed by a nonstem traversal, ending 3.62 ± 0.16 s and 10.29 ± 0.41 s after trial onset, respectively. This pulse was thus expected to suppress mPFC terminals in SuM throughout both traversals (Fig. 4C). After the experiments, we verified viral expression in mPFC, axonal labeling in SuM, and optic fiber placement (Fig. 4B, 4 animals).

We compared SuM and MS theta frequency between laser-off and laser-on trials. In laser-off trials, SuM theta frequency before the turn decreased during stem relative to nonstem traversals (stem 8.34 Hz, nonstem 8.47 Hz; 10 sessions from 4 rats; p < 0.001, linear mixed-effects model; Fig. 4D), replicating the result obtained in non-injected animals (Fig. 1). When mPFC terminals in SuM were suppressed, this modulation was absent (stem 8.19 Hz, nonstem 8.20 Hz; p = 1.000; Fig. 4D). MS theta frequency was unaffected by the suppression (control p = 0.965, laser p = 0.23).

These results show that mPFC input to SuM is required for the decision-dependent modulation of SuM theta frequency, identifying an active mechanism by which cortical input controls a subcortical theta oscillator.

### Theta-rhythm coordination between SuM and MS underlies decision-dependent mPFC– CA1 coupling

What is the wider network consequence of this change in frequency offset between SuM and MS theta? The reduced offset before the trajectory decision brings the two rhythms into a more stable phase relationship, which should in turn affect theta coordination among their downstream cortical targets.

We therefore examined mPFC and hippocampal CA1, which are coordinated by theta rhythm during trajectory decisions in spatial alternation tasks (*25, 28, 30*). Both SuM and MS project to mPFC and the hippocampus. SuM innervates primarily the dentate gyrus and CA2 (*54, 55*), whereas MS projects across hippocampal subregions (*56, 57*), yet SuM silencing alters mPFC– CA1 theta coordination (*30*), indicating that SuM can influence this coordination despite little direct projection to CA1.

We implanted LFP electrodes in mPFC, CA1, SuM, and MS to examine the relationship between SuM–MS frequency matching and mPFC–CA1 theta coordination. We first examined theta-band spectral coherence between mPFC and CA1 theta without laser application. We did not find a significant difference in the power of theta oscillations between stem and nonstem traversals before the turn (mPFC: p = 0.339; CA1: p = 0.266, Wilcoxon signed-rank tests). In contrast, mPFC–CA1 theta coherence was higher during stem than during nonstem traversals (stem: 0.46 ± 0.03; nonstem: 0.44 ± 0.03; mean ± s.e.m.; p = 0.013, repeated-measures ANOVA with animal identity as a factor), indicating that the coherence difference was not attributable to changes in theta power. We then analyzed the relationship between the SuM–MS frequency offset and mPFC–CA1 theta coherence and found a significant negative correlation (slope = −0.112 ± 0.015 per Hz, p < 0.001, linear mixed-effects model; Fig. 5B and 5D). This result suggests that frequency matching between SuM and MS is reflected in theta coordination between mPFC and CA1.

**Fig. 5.**
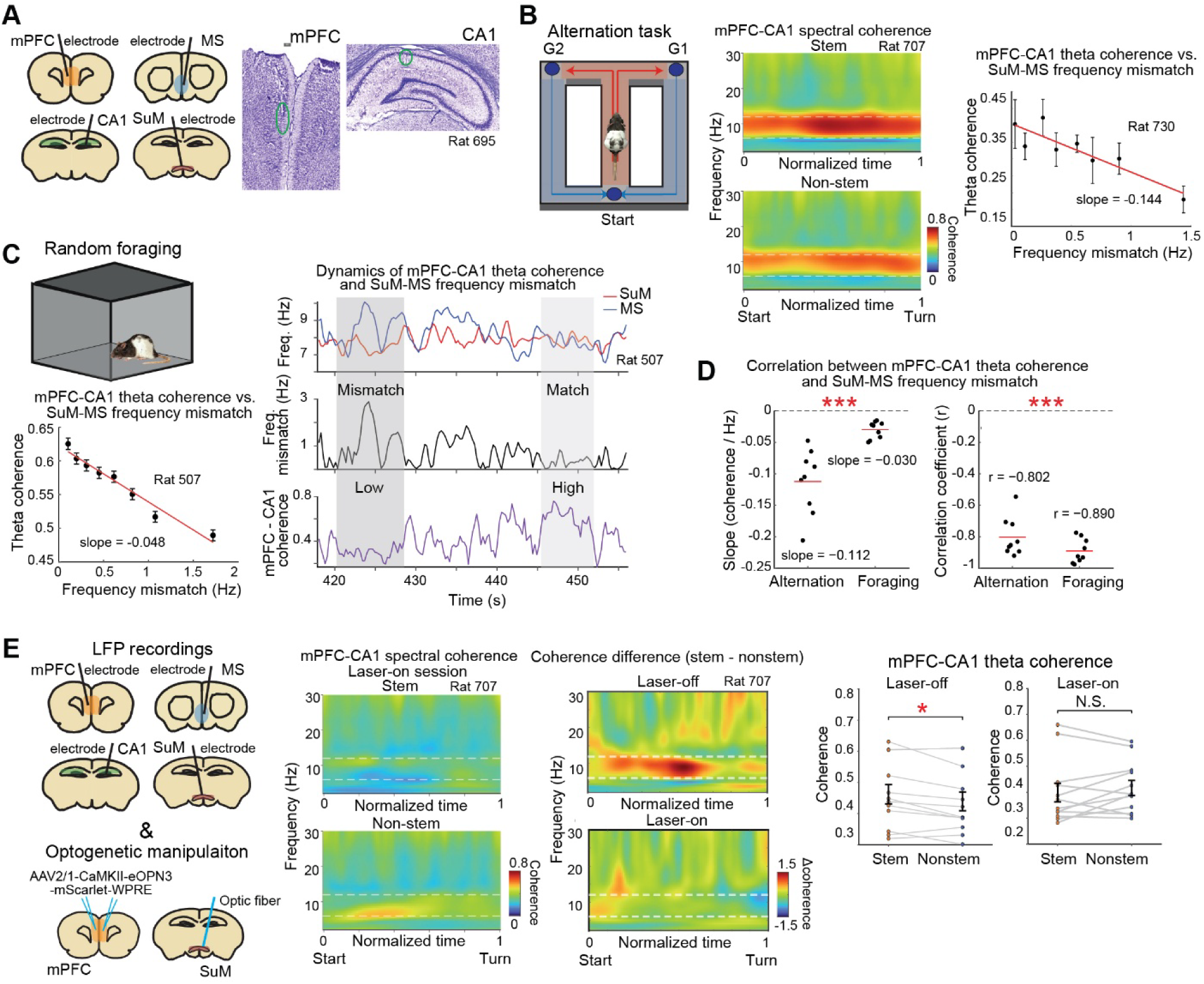
mPFC–CA1 theta coherence increases as SuM–MS frequency mismatch decreases and is disrupted by silencing of mPFC input to SuM. **(A)** Simultaneous LFP recordings from mPFC, MS, CA1, and SuM. Left, schematic of recording sites. Right, representative histological sections showing electrode tracks in mPFC and CA1. **(B)** Modified T-maze alternation task. Left, task schematic. Middle, mPFC–CA1 theta coherence is higher during stem than during nonstem traversals, as previously reported (*25, 28*). Right, mPFC– CA1 theta coherence as a function of SuM–MS theta-frequency mismatch. **(C)** Open-field random foraging, example session. Top left, arena (1 × 1 m). Right, from top to bottom: SuM and MS theta frequency, SuM–MS frequency mismatch, and mPFC–CA1 theta coherence. Periods of frequency match coincide with high coherence, and periods of mismatch coincide with low coherence. Bottom left, mPFC–CA1 theta coherence as a function of SuM–MS frequency mismatch (slope = −0.048/Hz; r = −0.977, p < 0.001). **(D)** Population summary. Regression slopes (left) and correlation coefficients (right) between SuM–MS frequency mismatch and mPFC–CA1 theta coherence for each session in both tasks. Slopes were negative in both tasks (p < 0.001, linear mixed-effects model with animal as random intercept), as were correlation coefficients (both p < 0.001). **(E)** Optogenetic silencing of mPFC input to SuM. Left, strategy for pathway-specific inhibition of mPFC axon terminals in SuM, with simultaneous LFP recordings from mPFC, MS, CA1, and SuM. Middle, example sessions: mPFC–CA1 theta coherence during stem and nonstem traversals with the laser on (left), and the stem minus nonstem coherence difference for laser-off and laser-on sessions (right). Right, population summary across sessions and animals. Coherence was higher during stem than nonstem traversals with the laser off (p = 0.013, repeated-measures ANOVA with animal identity as a factor) but not with the laser on (p = 0.346; 12 paired sessions, 2 rats).

We next asked whether this relationship extended beyond the spatial alternation task. We thus recorded while rats foraged randomly in an open-field arena (9 sessions from 3 rats; Fig. 5C). Theta oscillations were prominent during running, but mPFC–CA1 theta-band spectral coherence fluctuated over time. We estimated instantaneous theta frequency in SuM and MS as in the alternation task (Fig. 1), computed the frequency offset between them, and related it to mPFC–CA1 coherence. In a representative session, mPFC–CA1 theta coherence decreased as the SuM– MS frequency offset increased (linear regression slope = −0.048 per Hz; Pearson’s r = −0.98, p < 0.001; Fig. 5C). The same negative relationship held across sessions (slope = −0.030 per Hz, p < 0.001, linear mixed-effects model; 9 sessions from 3 rats; Fig. 5D). These results indicate that a reduced frequency offset between SuM and MS theta is associated with stronger theta coordination between their downstream targets, mPFC and CA1, in both the spatial alternation task and open-field foraging.

Given the influence of mPFC over SuM theta frequency and the SuM–MS frequency offset, we asked whether disrupting mPFC input to SuM would alter theta coordination between mPFC and CA1. We therefore recorded LFPs in mPFC and CA1 of rats expressing eOPN3 in mPFC, with an optic fiber positioned over SuM to silence mPFC terminals there (12 paired sessions from 2 rats; Fig. 5E). Without laser, mPFC–CA1 theta coherence was higher during stem than during nonstem traversals (reported above; Fig. 5E). In contrast, suppression of mPFC terminals in SuM abolished this segment-dependent enhancement (stem: 0.40 ± 0.04; nonstem: 0.42 ± 0.03; p = 0.346, repeated-measures ANOVA with animal identity as a factor). Theta power in mPFC and CA1 did not differ between control and suppression sessions (mPFC: p = 0.584, CA1: p = 0.428; repeated-measures ANOVA). Although running speed was reduced overall during the laser session (no laser: 78.60 ± 1.92 cm/s; laser: 67.58 ± 2.87 cm/s, p = 0.001, linear mixed-effects model), we did not observe a difference in speed between stem and nonstem traversals during either laser-off (p = 0.439) or laser-on (p = 0.885) sessions. Task performance was unaffected (error rate: no laser: 4.43 ± 0.91%; laser: 3.22 ± 0.93%, binomial generalized linear mixed-effects model, p = 0.133). Preserved performance was expected, as this task can be solved without an intact hippocampus (*58*). In the same task, however, SuM silencing disrupts mPFC–CA1 theta coordination and impairs the transfer of trajectory information from mPFC to CA1 (*30*), indicating that this coordination supports interregional information transfer even where it is not required for choice accuracy. The loss of mPFC–CA1 theta coordination therefore followed the loss of decision-dependent modulation of the SuM–MS frequency offset, without accompanying changes in task performance, local theta power, or the segment-dependent structure of locomotion. These results are consistent with mPFC input to SuM reorganizing temporal coordination between distant cortical regions through frequency matching of subcortical oscillators, without altering the local oscillations themselves.

Together, these results link mPFC-dependent modulation of the SuM–MS frequency offset to theta coordination between mPFC and CA1. The closer the two subcortical rhythms were in frequency, the stronger mPFC–CA1 theta coherence became. These findings identify an active control mechanism for interregional coupling, in which independently tunable subcortical oscillators allow cortical coordination to be adjusted according to behavioral demand.

## Discussion

Coordinated oscillatory activity has long been proposed to provide a temporal framework for selective interaction and information routing between distributed neural populations (*2–8*), yet whether and how cortical synchrony is actively controlled in accordance with behavioral demands has remained unknown. Here we identify a circuit mechanism by which this temporal organization is reconfigured by top-down prefrontal control. As rats ran toward the trajectory choice point, SuM theta frequency decreased while MS theta frequency remained stable, reducing the frequency mismatch between the two oscillators. The same dissociation was present in population spiking. Immunostaining for synaptic markers identified putative direct projections from mPFC to SuM, and optogenetic suppression of this input abolished the decision-dependent modulation of SuM theta frequency. This offset tracked theta coordination between the downstream targets of the two oscillators, mPFC and CA1: theta coherence was stronger when the two frequencies were closely matched and weaker as they diverged. This relationship was observed in both the spatial alternation task and open-field foraging, indicating a general mechanism for mPFC-CA1 coordination across behaviors. Finally, suppressing mPFC input to SuM abolished the segment-dependent enhancement of mPFC-CA1 theta coherence, consistent with prefrontal control of the SuM-MS frequency offset underlying behavior-dependent cortical theta coupling.

A central feature of brain computation is that information is processed locally and in parallel. In the visual system, individual regions process distinct parts of the visual field. Such parallel processing is efficient, but it raises the question of how separately processed features are integrated into a coherent percept, the so-called binding problem (*59*). Oscillatory synchrony was proposed as a solution: neurons representing features of the same object fire in a common temporal window, which binds them without changing anatomical convergence (*1, 60*). The idea was subsequently extended from local circuits to interactions between regions, each with a specialized role (*12*). Many studies have since reported that interregional synchrony changes with behavioral demand (*7, 9, 11, 25, 28*). These findings are largely correlational, however, leaving open whether synchrony regulates neural communication or reflects the underlying network activity. Experimental and modeling work has shown that interregional oscillatory coherence can emerge from synaptic transmission and vary with oscillatory power and anatomical connectivity, without requiring phase synchronization between local oscillators (*61–63*), implying that synchrony may be a consequence of communication rather than a mechanism that controls it. Whether the brain possesses an active mechanism for setting oscillatory coherence is thus central to this debate.

Theta oscillations are a dominant form of activity in the hippocampal–entorhinal system and have been implicated in multiple forms of information processing: they gate synaptic plasticity (*64*), organize spatial coding (*17*), and support associative memory (*50*). How theta is generated and controlled has therefore been a long-standing question. Recordings in anesthetized animals identified two subcortical theta oscillators, SuM and MS. Neurons in both regions fire theta-rhythmically, but procaine injection into SuM reduced theta frequency, whereas the same injection into MS reduced mainly theta power (*39–43, 65*), placing SuM in control of hippocampal theta frequency. Awake animals gave a different picture: silencing MS nearly abolished hippocampal theta, whereas SuM lesions did not (*35, 41–43, 66*). MS was therefore regarded as the pacemaker and SuM as its modulator. The anesthetized results, however, indicate that SuM can express theta rhythmicity on its own. Consistent with this, SuM lesions reduce hippocampal theta frequency while leaving theta itself intact (*43, 67*), and rhythmic optogenetic stimulation of SuM drives hippocampal theta (*68*). Here we show that SuM operates as an independently tunable oscillator whose frequency relationship with MS changes with behavioral demand. By identifying the prefrontal input that controls SuM theta frequency, we define a mechanism by which the brain can actively control cortical synchrony. For theta rhythm at least, this argues that oscillatory synchrony is not a byproduct of network activity but a variable the brain sets to organize cortical communication.

We found that mPFC axons innervated SuM, and that presynaptic and postsynaptic markers colocalized along these axons, indicating putative synaptic contacts onto SuM neurons. Optogenetic silencing of mPFC terminals in SuM abolished the behavior-dependent shift in SuM rhythmic frequency, showing that the mPFC–SuM projection is functional and necessary for the shift. At the single-neuron level, 126 of 140 SuM neurons with significant theta rhythmicity (maximum-likelihood estimation; Methods) were rhythmic in only one segment. This raised the possibility that behavioral demand recruits distinct subsets of neurons, each preferring a different frequency. Two observations argue against this interpretation. First, the neurons rhythmic in both segments showed lower rhythmic frequency during stem than nonstem traversals. Second, because autocorrelation-based estimates require many spikes, segment-specific rhythmicity could instead reflect segment-dependent differences in spike count. We therefore computed population-level frequency-tuning profiles within the theta band (6–10 Hz) from the phase relationships of within-neuron spike pairs, which allowed us to include every recorded neuron. During stem traversals, the SuM population frequency profile shifted toward lower frequencies, with no detectable change in peak width and no evidence of multimodality. This suggests that the frequency shift was population-wide rather than driven solely by recruitment of distinct frequency-specific subsets. How mPFC input alters the rhythmic frequency of SuM neurons remains an open question, but two mechanisms are plausible. First, the effect could be synaptic. Changes in direct excitation, or in feedforward inhibition through local SuM interneurons, would alter the net drive onto rhythmic SuM neurons and change their spiking frequency. Second, the effect could be neuromodulatory. SuM neurons are heterogeneous in their expression of neuropeptides and neuromodulators (*68–70*), and mPFC input could trigger their local release. These signals could then modulate conductance that sets membrane resonance in rhythmic SuM neurons and shift their preferred frequency.

Silencing mPFC input to SuM abolished the behavior-dependent shift in SuM theta frequency and disrupted mPFC–CA1 theta coordination during stem traversals. This coordination has a functional consequence. During stem traversals, mPFC neurons encode the upcoming trajectory (left or right turn). The nucleus reuniens relays this signal to CA1, where place cells express it as trajectory-dependent “splitter” firing (*31, 71, 72*). Silencing SuM itself disrupts the decision-dependent increase in mPFC–CA1 theta coordination and reduces trajectory coding in CA1 (*30*). Here, selectively silencing mPFC input to SuM reproduced the loss of coordination. Task performance was unaffected, as expected because this task does not require an intact hippocampus (*58*). The loss of mPFC–CA1 coordination therefore reflected the circuit manipulation rather than a secondary consequence of altered behavior.

Recordings during open-field foraging suggest that SuM–MS frequency matching is a general mechanism for cortical theta coupling. As in the spatial alternation task, mPFC–CA1 coordination was enhanced when SuM and MS frequencies were closely matched and disrupted when they diverged. During foraging, however, the mismatch fluctuated continuously and did not follow segment-specific behavior as in the spatial alternation task. Moreover, MS theta frequency was on average lower than SuM frequency in the alternation task but higher during foraging. Despite this reversal, the magnitude of the mismatch had the same effect on mPFC–CA1 coordination in both conditions. This indicates that the frequency difference between the two oscillators, rather than their absolute frequency, governs cortical theta coupling.

Coupling through two independently tunable oscillators could extend to other frequency bands and brain regions. It does, however, place a requirement on connectivity: target regions must differ in their relative input from the two oscillators. mPFC and CA1 both receive direct or indirect input from SuM and MS (*30, 54–57, 73*), presumably in different proportions. If both regions were dominated by the same oscillator, their coupling would be insensitive to frequency mismatch. Conversely, coupling would be most sensitive to mismatch if each region were driven by a different oscillator. More likely, individual regions receive graded input from both, so that each region’s entrainment depends on its input balance. Small mismatch would couple regions broadly, whereas large mismatch would restrict coupling to regions with similar input balance, providing a mechanism for selective, dynamic routing of information across distributed networks. The capacity to control this mismatch would expand the brain’s computational repertoire, reconfiguring information flow according to behavioral demand without changes in anatomical connectivity and providing a potential substrate for parallel computation.

## Methods

### Animals

Experiments were performed on 17 adult male Long–Evans rats, aged 3–6 months and weighing 400–550 g at the start of the experiments. All procedures were approved by the Regierungspräsidium Darmstadt, Germany (protocols F126/1009, F126/1026 and F126/2005) and by the veterinary authority of the Canton of Vaud, Switzerland (protocol VD4053). Rats were housed individually in enriched Plexiglas cages (45 × 35 × 40 cm; Tecniplast GR1800) on a reversed 12-h light/dark cycle, and all behavioral experiments were conducted during the dark phase. Rats had unrestricted access to food and were water-restricted, with body weight maintained at 85–90% of the pre-restriction value; body weight and welfare were monitored daily. Four rats were implanted with LFP electrodes in SuM and MS. One rat received a tetrode microdrive targeting SuM and MS together with LFP electrodes in mPFC and CA1, and performed both the alternation task and open-field foraging. Two rats underwent four-region LFP recordings (SuM, MS, mPFC, and CA1) during open-field foraging. Two rats received four-region LFP implants together with AAV-CamKIIα-eOPN3-mScarlet-WPRE (Addgene #125712) injection in mPFC and an optic fiber targeting SuM, and performed the alternation task. Two rats were implanted with two Neuropixels 2.0 probes and one rat with two Neuropixels 1.0 probes for simultaneous SuM and MS recordings. Two rats received a microdrive combining tetrodes and an optic fiber (optrode) in SuM and MS together with LFP electrodes in CA1. Three rats were used for anatomical tracing.

### Surgery, viral injection, and electrode implantation

All surgical procedures were performed under isoflurane anesthesia in a stereotaxic frame (David Kopf Instruments). Anesthesia was induced with 5% isoflurane and maintained at 0.5–2%, adjusted according to physiological monitoring. Analgesia was provided by subcutaneous buprenorphine (Buprenovet, 0.06 mg/mL, WdT; or Temgesic, 0.1 mg/kg), together with subcutaneous injection of bupivacaine (0.5 mg/mL, Jenapharm), ropivacaine (2 mg/mL, Fresenius Kabi), or a combination of lidocaine (6 mg/kg) and bupivacaine (2.5 mg/kg), into the scalp. Anesthetic depth and physiological condition were monitored throughout surgery, and body temperature was maintained with a heating pad. After induction, the rat was secured in the stereotaxic frame, the scalp was disinfected and incised, and the skull was exposed and leveled. Small openings were drilled for anchor screws, followed by craniotomies above the targeted recording or injection sites. Microdrives, electrodes, and optical fibers were secured to the skull with anchor screws and dental cement. Postoperatively, rats received meloxicam (Metacam, 2 mg/mL; Boehringer Ingelheim) and the antibiotic enrofloxacin (Baytril, 25 mg/mL; Bayer), or paracetamol (1500 mg/800 mL) and carprofen (40 mg/800 mL) in drinking water. Rats were monitored daily during recovery and were allowed at least 7 days before the start of recordings.

Tetrodes were fabricated from 17-µm polyimide-coated platinum–iridium wires (90:10%; California Fine Wire) and gold-plated to reduce impedances below 150 kΩ at 1 kHz. The tetrodes were assembled into bundles using stainless-steel guide cannulas and mounted on adjustable microdrives, allowing individual electrodes to be advanced toward the recording targets. Each microdrive carried two bundles, targeting SuM and MS for simultaneous recording. LFP electrodes were constructed from the same tetrode wire, with all four channels shorted together to yield a single signal. Two skull screws positioned above the cerebellum were connected to the recording-system ground. For simultaneous high-density recordings from SuM and MS, rats were implanted with two Neuropixels 1.0 or 2.0 probes.

Stereotaxic implantation coordinates relative to bregma were as follows: mPFC: anterior–posterior (AP) +3.25 mm, mediolateral (ML) 0.5 mm, dorsoventral (DV) 3.0 mm, with a 4° implantation angle; hippocampal CA1: AP −4.0 mm, ML 3.5 mm, DV 2.5 mm; MS: AP +0.5 mm, DV 6.0 mm, with a 5° angle toward the midline on the coronal plane; and SuM: AP −4.5 mm, DV 8.0 mm, with a 5° angle toward the midline on the coronal plane. Electrodes were positioned according to the targeted anatomical structures and subsequently adjusted to optimize neural recordings.

For anatomical tracing, an adeno-associated viral (AAV) vector carrying the SYN1–mApple– WPRE expression cassette was stereotaxically injected into mPFC across its dorsoventral extent to label axonal projections to SuM. These rats underwent tracing only, without electrophysiological implantation or optogenetic manipulation. For pathway-specific optogenetic inhibition, an AAV encoding the inhibitory opsin eOPN3 (*53*) under the CaMKIIα promoter was injected into mPFC. Injections were performed bilaterally with a microsyringe connected to an infusion pump at 100 nl/min, at the following coordinates (mm from bregma): AP +3 and +3.75, ML ±0.5, DV 4.5, 3.5 and 2.5 from the brain surface, with a 4° angle toward the midline on the coronal plane. The needle was left in place for 5 min after each infusion to minimize reflux before withdrawal. Optical fibers (FP400URT, Thorlabs) were implanted approximately 500 µm above SuM for delivery of 532-nm laser light from a DPSS laser (Shanghai Laser & Optics Century). Movable tetrodes were advanced toward their targets over the following 2–4 weeks, and optogenetic experiments began at least four weeks after viral injection to allow transgene expression. After completion of the experiments, electrode tracks, Neuropixels probe locations, optical fiber positions, and viral expression were verified histologically.

### Histological procedures and immunohistochemistry

Following completion of the experiments, animals were deeply anesthetized with sodium pentobarbital and perfused transcardially with saline followed by 10% formalin. Brains were extracted and post-fixed in formalin for at least 72 h at 4°C. For anatomical verification of implanted electrodes, optical fibers, and Neuropixels probes, coronal sections (30 µm) were prepared using a cryostat, mounted onto glass slides, stained with cresyl violet, and coverslipped. Electrode trajectories and implantation sites were identified by examining consecutive sections. For anatomical tracing of mPFC projections, three male rats received injections of AAV-Syn1-mApple-WPRE. Free-floating coronal sections (50 µm) were prepared for immunohistochemistry. To enhance the fluorescent reporter signal, sections were washed in phosphate-buffered saline (PBS) for 50 min and blocked for 1 h at room temperature in PBS containing 10% fetal bovine serum and 0.1% Triton X-100. Sections were incubated overnight at 4°C with rabbit anti-RFP antibody (1:500; Rockland, 600-401-379), followed by four 5-min PBS washes. Alexa Fluor 594-conjugated anti-rabbit secondary antibody (1:500; Abcam, ab150080) and Hoechst (1:2000) were applied for 1.5 h at room temperature. Sections were subsequently washed three times in PBS and mounted on Superfrost slides using Fluoroshield mounting medium (Agilent Technologies).

To identify putative synaptic contacts formed by mPFC axons in SuM, additional sections were triple-immunostained using guinea pig anti-Bassoon (1:1000; Synaptic Systems, 141 318), mouse anti-PSD-95 (1:600; Merck, MAB1596, clone 6G6-1C9), and rabbit anti-RFP (1:500; Rockland, 600-401-379). Following overnight primary-antibody incubation at 4°C, sections were incubated with Alexa Fluor 647-conjugated anti-guinea pig (1:1000; Abcam, ab150187), Alexa Fluor 488-conjugated anti-mouse (1:1000; Abcam, ab150113), and Alexa Fluor 594-conjugated anti-rabbit (1:500; Abcam, ab150080) secondary antibodies, together with Hoechst (1:2000). Sections were mounted using Fluoroshield mounting medium. Fluorescently labeled axons and the spatial association of Bassoon-positive presynaptic puncta with PSD-95-positive postsynaptic puncta were examined by confocal microscopy.

For optogenetic experiments, mScarlet fluorescence associated with eOPN3 was examined to confirm expression in mPFC and labeling of axonal projections in SuM, using the anti-RFP immunostaining described above, and optical fiber placement was verified in cresyl violet-stained sections.

### Behavioral training, spatial alternation task, and open-field foraging

Rats were habituated to the experimental apparatus and trained to perform a continuous spatial alternation task on a modified T-maze (1 × 1 m). The maze comprised a central stem, a start well, two goal wells, and return pathways arranged so that outward and return trajectories were geometrically matched. On each trial, the rat ran from the start well along the central stem and entered either the left or right arm. The rewarded arm alternated between trials, and correct choices were reinforced with a water reward at the goal well. After reward consumption, the rat returned to the start well along the corresponding return pathway, consuming a reward and initiating the next trial. Training proceeded in stages. Rats were first habituated to the maze with water available at all wells. They were then trained to run in one direction along the maze, with both arms rewarded. Finally, the alternation rule was introduced, with reward delivered only at the arm opposite to the previously visited one. Rats were considered proficient when they made correct choices on at least 90% of trials. For analysis, journeys from the start well to a goal well, which required a choice of turn direction, were designated stem trials; journeys from a goal well back to the start well along the side arms were designated nonstem trials. Each trial contained a single turn, and the segments preceding and following it were analyzed separately as the pre-turn and post-turn phases. Stem and nonstem traversals were compared over geometrically corresponding portions of the two trajectories. Position and head direction were tracked from two LEDs on the head implant, recorded by an overhead camera at 20 Hz, and used to delineate journey segments and turning points and to compute running speed and acceleration.

For open-field recordings, rats foraged for cookie crumbs scattered on the floor of a 1 × 1 m arena. This condition provided sustained locomotion without the structured trajectory decisions imposed by the maze.

### Recording procedures and spike sorting

Electrophysiological recordings were obtained from freely moving rats using chronically implanted tetrodes, Neuropixels 1.0 and 2.0 probes (*74, 75*), and LFP electrodes. Signals from tetrode microdrives were acquired and amplified with RHD2164 headstages (Intan Technologies) at 15 kHz using the Open Ephys acquisition system, and LFP signals with RHD2132 headstages (Intan Technologies) on the same system. Neuropixels signals were acquired with a PXIe acquisition module (IMEC) housed in an NI PXIe-1071 chassis (National Instruments), also using Open Ephys. Signals were band-pass filtered between 0.6 and 6 kHz for spike detection. LFP signals from SuM, MS, mPFC and CA1 were sampled or downsampled to 1 kHz for theta-frequency and coherence analyses.

Spike detection and clustering were performed using Kilosort (https://github.com/cortex-lab/KiloSort) (*76*). Tetrode recordings were sorted with Kilosort 1, with tetrode-specific channels grouped through the ‘kcoords’ parameter and the shared-noise parameter set to 0.01. Automatically detected clusters were then inspected using spike autocorrelograms, waveform characteristics and separation in principal-component feature space, and noise clusters and poorly isolated multiunit activity were discarded to obtain well-isolated single units. Neuropixels recordings were sorted with Kilosort 3, and individual units were assigned to SuM or MS from their recording-channel positions together with histological reconstruction of the probe trajectories. Spike times were aligned to simultaneously acquired LFP signals, LED position tracking, and maze event timestamps. Spike trains were then segmented into stem and nonstem epochs for analyses of rhythmic firing, autocorrelation structure, and spike-field coherence.

### Optogenetic inhibition

For pathway-specific suppression of medial prefrontal cortex (mPFC) projections to the supramammillary nucleus (SuM), the Gi/o-coupled bistable rhodopsin eOPN3 (*53*) was expressed in mPFC neurons using AAV2/1-CaMKIIα(0.4)-eOPN3-mScarlet (Addgene #125712). Illumination of eOPN3-expressing terminals suppresses presynaptic neurotransmitter release through Gi/o signaling, with suppression reaching saturation within ∼1 s of light onset and recovering spontaneously over several minutes (*53*).

An optical fiber was implanted approximately 500 µm dorsal to SuM (AP -4.5 mm, ML 0.5 mm, DV 7.5 mm with a 5° angle toward the midline on the coronal plane) to illuminate eOPN3-expressing mPFC axon terminals. In these animals, LFPs were recorded simultaneously from SuM, MS, mPFC and CA1.

Optogenetic experiments began at least four weeks after viral injection to allow transgene expression. Green light (532 nm) was delivered from a DPSS laser (Shanghai Laser & Optics Century) with output adjusted to 10 mW at the fiber tip. During the continuous alternation task, a single 500-ms pulse was delivered on each trial while the rat consumed reward at the start well, before journey onset. Because eOPN3-mediated suppression saturates within ∼1 s and decays over several minutes (*53*), one pulse per trial maintained terminal silencing at near steady state throughout the session while confining illumination to stationary epochs, avoiding photoelectric artifacts during stem and nonstem traversals.

Laser-off and laser-on sessions were run sequentially on the same day. Theta frequencies in SuM and MS, mPFC–CA1 theta coherence, running speed and task performance were compared between conditions. After completion of the experiments, viral expression in mPFC, labeled axonal projections in SuM, and optical fiber placement were verified histologically.

### LFP theta-frequency estimation and validation

LFPs recorded simultaneously from SuM and MS were analyzed to estimate instantaneous theta frequency. Signals were sampled at 1 kHz and band-pass filtered between 6 and 12 Hz with a fourth-order Butterworth filter applied forward and backward to avoid phase distortion. The analytic signal was obtained by Hilbert transform, and the instantaneous phase was unwrapped. Instantaneous frequency was computed as the temporal derivative of the unwrapped phase divided by 2π. Frequency estimates outside the 4–14 Hz range were excluded. Frequency traces were then median filtered over a 100-ms sliding window to suppress brief, spurious frequency fluctuations arising from noise-sensitive phase differentiation and smoothed with a 100-ms moving average, both applied along time within each traversal. Frequency estimates were aligned to the normalized behavioral trajectories. The instantaneous frequency difference between the two subcortical regions was calculated as Δf(t) = fSuM(t) − fMS(t), preserving the sign of the offset. Mean theta frequencies and SuM–MS frequency differences were calculated separately for stem and nonstem traversals in the pre-turn and post-turn phases.

Time-resolved power spectra were computed by short-time Fourier transform over 3–12 Hz using a 0.75-s window advanced in 0.05-s steps. Each spectrogram was normalized by dividing the power at each frequency by its session average. For visualization, trial-averaged power spectra were converted to decibels and normalized by subtracting the median and dividing by the median absolute deviation across the combined stem and nonstem time–frequency representation.

Frequency estimates were validated with a second method, the synchrosqueezed wavelet transform (*48*). Synchrosqueezing reassigns wavelet coefficients toward the instantaneous-frequency ridge, sharpening the time–frequency representation and yielding frequency estimates without the narrow-band prefiltering required by the Hilbert approach. SuM and MS estimates were extracted over the same behavioral epochs as in the Hilbert analysis, and condition-dependent frequency differences were reassessed with the same linear mixed-effects models.

The reliability of behavioral frequency modulation was further assessed by randomly splitting stem and nonstem trials into two halves and comparing the SuM–MS frequency offset between halves of the same condition and between halves of different conditions. To test how this offset constrains oscillatory coordination, pairs of sinusoidal oscillators were simulated across the observed range of frequency offsets, and their phase coherence was computed over 1.62 s, the mean duration of the pre-turn trajectory, yielding coherence as a function of frequency offset.

### Single-unit rhythmicity, autocorrelation, and frequency estimation

Spike-train rhythmicity was analyzed separately for SuM and MS neurons during stem and nonstem traversals, following the model-based approach of Climer et al. (*49*), which separates oscillatory spike timing from non-rhythmic autocorrelation structure. Spike trains were segmented by trial and binned at 5 ms. For each trial, the binned train was mean-centered and its normalized autocorrelation computed for lags up to 500 ms. Trials shorter than 1.4 s or containing fewer than two spikes were excluded.

Intrinsic rhythmic frequency was estimated by fitting the autocorrelation over positive lags of 50– 500 ms. A non-oscillatory model, comprising a constant baseline and two exponential decay components, was compared with models that added a damped sinusoidal component, evaluated over candidate frequencies of 5–10 Hz and a predefined range of decay constants. We selected models using the Bayesian information criterion (BIC), and we took the rhythmic frequency from the best-fitting oscillatory model.

Evidence for rhythmicity was assessed by trial-preserving spike-time shuffling. Surrogate trains were generated by randomly redistributing spike times within each trial, preserving trial duration and spike count. A neuron was classified as theta-rhythmic when its rhythmicity score exceeded the surrogate criterion, the condition-specific shuffle probability was < 0.05, and the fitted frequency fell strictly within 5–10 Hz. Classification was performed independently for stem and nonstem traversals; neurons rhythmic in at least one condition were termed either-rhythmic and those rhythmic in both, both-rhythmic.

For visualization, within-trial spike lags were pooled across trials of the same condition, without spike pairs spanning trial boundaries. The pooled autocorrelations were fitted with a phase-constrained damped cosine, the preferred frequency selected by BIC over 5–10 Hz. These fits served only to display autocorrelation peaks and their temporal progression; the rhythmicity classifications above were unchanged. Frequency distributions were compared between stem and nonstem traversals by two-sample Kolmogorov-Smirnov tests. For neurons rhythmic in both conditions, frequencies were compared with a linear mixed-effects model with condition as a fixed effect and animal, session, and neuron as nested random effects.

To assess the stability of rhythmic-frequency estimates, stem and nonstem trials were each divided chronologically into first and second halves, with at least three trials and 15 spikes required per half. Autocorrelations and rhythmic frequencies were recomputed for each half; neuron classification was held fixed from the full-session fits. The two within-condition frequency differences (first versus second half of the same condition) were then tested against the four cross-half stem-nonstem differences. Cross-half differences were averaged within each neuron beforehand, so that repeated measurements were not treated as independent, and comparisons used two-sided Wilcoxon signed-rank tests.

### Spike–field coherence and local theta phase locking

To relate neuronal rhythmic firing to local theta oscillations, spike–field coherence was computed for neurons classified as rhythmic in both stem and nonstem traversals, using spikes and simultaneously recorded LFPs from SuM and MS. Spike times were binned at 1 ms, and LFP segments of ±250 ms were extracted around each spike. Spike–field coherence was then computed as described previously (*50, 51*): the power spectrum of the spike-triggered average LFP was divided by the mean power spectrum of the contributing LFP segments, each spectrum obtained with a single Hann taper. The ratio approaches unity at frequencies where the segments are phase-aligned and falls toward zero where spike timing is unrelated to LFP phase. To equate trial numbers across conditions, 80% of the smaller of the available stem and nonstem trial counts was drawn at random on each of 200 iterations, and coherence spectra were averaged over iterations. Neurons were included when at least four valid trials and 20 spikes were available per condition. The dominant coupling frequency was taken as the coherence maximum within 6-10 Hz, refined by quadratic interpolation about the spectral peak. Stem and nonstem peak frequencies were compared with a linear mixed-effects model with condition as a fixed effect and animal, session, and neuron as nested random intercepts. Peak coherence over 6–10 Hz was analyzed separately to test whether shifts in peak frequency were accompanied by changes in coupling strength.

Theta spike-phase locking was examined with a second phase-based measure. LFPs were filtered between 6 and 10 Hz, and instantaneous phase and amplitude were extracted with the Hilbert transform. LFP phase was sampled at each spike time, and neurons with fewer than 20 valid phase samples in either condition were excluded. Phase-locking strength was quantified as the mean vector length (MVL) of the circular spike-phase distribution. To limit spike-count-dependent bias, spike counts were equalized across conditions by random subsampling over 500 iterations, and the resulting MVL estimates were averaged. Phase locking was tested for significance with Rayleigh tests.

### Population spike-pair phase consistency and bootstrap analysis

To estimate rhythmic frequency from population spiking without relying on single-neuron rhythmicity classification, we quantified spike-pair phase consistency across candidate theta frequencies using all simultaneously recorded SuM and MS neurons. For each neuron and trial, we computed the intervals between all within-neuron spike pairs. At each candidate frequency *f*, the phase difference corresponding to a spike-pair interval Δ*t* was calculated relative to a reference sinusoid as Δφ = 2πfΔt, and the cosine of this phase difference was used to quantify rhythmic timing consistency. Neurons with fewer than two spikes in a trial were excluded from that trial. Candidate frequencies ranged from 6 to 10 Hz in 0.1-Hz steps. For each trial, cosine values from all valid within-neuron spike pairs were pooled across contributing neurons and divided by the total number of valid pairs, yielding a population phase-consistency profile across candidate frequencies. Each trial’s profile was then min–max normalized across frequencies. The population frequency profile for each behavioral condition was obtained by averaging normalized trial-level profiles and is displayed as the mean ± s.e.m. across trials. Peak frequency was defined as the candidate frequency at the maximum of the trial-averaged profile, without curve fitting or interpolation. Stem and nonstem trials were analyzed separately. Differences in peak frequency between behavioral conditions were tested independently for SuM and MS using linear mixed-effects models, with behavioral condition as a fixed effect and session nested within animal as random intercepts.

To determine whether the behavioral modulation of population rhythmic frequency was accompanied by changes in the shape of the frequency profile, we examined peak width and modality. Peak width was quantified using the full width at half maximum (FWHM) and compared between stem and nonstem conditions using linear mixed-effects models. Modality was assessed using Hartigan’s dip test applied to the distribution of trial-wise peak frequencies pooled across sessions, with significance determined by bootstrap resampling under a uniform null distribution.

To control for differences in the numbers of simultaneously recorded SuM and MS neurons, we performed an equal-neuron-count bootstrap analysis. Within each recording session, neurons were resampled with replacement from both regions, using the smaller regional population to determine the number of neurons sampled from each. Population frequency estimates were reconstructed separately for stem and nonstem trials, and 200 pseudo-trials were generated per iteration to estimate the behavioral frequency shift within each region and the difference in frequency shifts between regions. This procedure was repeated 3,000 times separately for each rat. Regional contrasts were averaged across rats within each iteration to prevent animals contributing more recording sessions from disproportionately influencing the results. The resulting rat-balanced distributions were summarized by their median and 95% percentile intervals. Directional significance was estimated as the proportion of bootstrap iterations yielding a contrast with the opposite sign to the observed effect.

### Analysis of optogenetic manipulation

To test whether mPFC input to SuM contributes to the behavioral modulation of theta frequency, LFPs were analyzed during pathway-specific inhibition of mPFC terminals in SuM. Recordings were obtained from rats expressing eOPN3 in mPFC neurons, with simultaneous LFPs from SuM and MS. Trials were classified by behavioral condition (stem or nonstem) and optogenetic condition (laser-on or laser-off). Signals were aligned to normalized behavioral trajectories, and pre-turn and post-turn periods were analyzed separately, as described above.

Instantaneous theta frequency was estimated from SuM and MS LFPs with the Hilbert-based procedure described above. For each session, frequency estimates were averaged across trials within each behavioral segment and optogenetic condition. Stem and nonstem frequencies were then compared separately for laser-off and laser-on trials, and condition-dependent differences were tested with linear mixed-effects models with session nested within animal as random intercepts. SuM and MS were analyzed separately to test whether suppression selectively altered SuM frequency modulation, and the SuM–MS frequency difference was analyzed to characterize changes in the coupling between the two oscillators.

To assess behavioral consequences of optogenetic inhibition, running speed and task performance were quantified for laser-on and laser-off trials. Running speed was computed over the same epochs used for the electrophysiological analysis and compared with linear mixed-effects models. Performance was expressed as the proportion of incorrect alternation choices, and laser-associated differences in error probability were tested with a binomial generalized linear mixed-effects model. Both models included session nested within animal as random intercepts.

### mPFC–CA1 theta coherence and open-field analysis

To examine theta coordination between mPFC and CA1, simultaneously recorded LFPs were analyzed during the spatial alternation task and open-field foraging. For the alternation task, theta-band coherence between mPFC and CA1 was computed separately for stem and nonstem traversals under laser-off and laser-on conditions. Frequency-resolved coherence was aligned to the normalized pre-turn trajectory, and stem-minus-nonstem differences were computed across time and frequency. Mean theta-band coherence was then obtained for each session and condition, and stem and nonstem traversals were compared with repeated-measures ANOVA across 12 paired sessions from two rats. To test whether coherence differences were accompanied by changes in local oscillatory activity, theta power in mPFC and CA1 was analyzed with paired Wilcoxon signed-rank tests, and, for comparisons involving optogenetic inhibition, with repeated-measures ANOVA including animal identity as a factor.

To examine the relationship between subcortical theta-frequency matching and cortical coordination, SuM–MS frequency mismatch and mPFC–CA1 theta coordination were compared across the spatial alternation task and open-field foraging, with LFPs recorded simultaneously from all four regions. mPFC and CA1 signals were sampled at 1 kHz and band-pass filtered between 6 and 12 Hz with a zero-phase filter. Instantaneous phase was extracted with the Hilbert transform. The spectral coherence was calculated from the LFP cross-spectral and autospectral estimates.

Instantaneous theta frequency in SuM and MS was estimated with the Hilbert-based procedure described above, and the frequency mismatch computed as |fSuM(t) − fMS(t)|. Within each session, mPFC–CA1 coherence was regressed on simultaneous frequency mismatch, and the resulting slopes were tested across sessions with a linear mixed-effects model with session nested within animal as a random intercept, to test whether larger subcortical frequency mismatch was consistently associated with weaker cortical theta coordination.

### Statistical analysis

Statistical analyses were performed in MATLAB (MathWorks). Data are reported as mean ± s.e.m. unless otherwise indicated. The statistical unit (neuron, recording session, or animal) was determined by the design of each analysis and is stated with the corresponding result. Linear mixed-effects models were used wherever multiple sessions or neurons from the same animals contributed to a comparison, to account for the hierarchical structure of the recordings.

For LFP theta-frequency analyses, behavioral condition (stem versus nonstem), running speed and absolute acceleration were fixed effects, with session nested within animal as random intercepts. Time-resolved analyses additionally included signed acceleration and condition-specific session effects. For single-unit rhythmic frequency and spike–field coherence, behavioral condition was the fixed effect, with session and neuron nested within animal as random intercepts. Fixed-effect estimates are reported with 95% confidence intervals and p values.

Distributions of rhythmic frequency were compared with two-sample Kolmogorov–Smirnov tests. Phase locking was tested with Rayleigh tests, and uncertainty in circular phase differences was estimated by bootstrap resampling. Population frequency estimates were assessed with rat-balanced bootstrap procedures, with confidence intervals taken from the resulting distributions.

For optogenetic experiments, theta-frequency modulation was tested with linear mixed-effects models and behavioral error probability with a binomial generalized linear mixed-effects model. Theta power across optogenetic conditions was compared by repeated-measures ANOVA. The relationship between SuM–MS frequency mismatch and mPFC–CA1 theta coherence was assessed by linear regression within sessions and a linear mixed-effects model across sessions.

Tests were two-sided unless otherwise specified. Directional bootstrap probabilities were computed as the proportion of resampled estimates with the opposite sign to the observed effect. Numbers of animals, recording sessions and neurons are given for each analysis and figure panel.

## References

1. C. M. Gray, P. Konig, A. K. Engel, W. Singer, Oscillatory responses in cat visual cortex exhibit inter-columnar synchronization which reflects global stimulus properties. Nature 338, 334–337 (1989).

2. W. Singer, Neuronal synchrony: a versatile code for the definition of relations? Neuron 24, 49–65, 111–125 (1999).

3. A. K. Engel, P. Fries, W. Singer, Dynamic predictions: oscillations and synchrony in top-down processing. Nat Rev Neurosci 2, 704–716 (2001).

4. F. Varela, J. P. Lachaux, E. Rodriguez, J. Martinerie, The brainweb: phase synchronization and large-scale integration. Nat Rev Neurosci 2, 229–239 (2001).

5. G. Buzsaki, A. Draguhn, Neuronal oscillations in cortical networks. Science 304, 1926–1929 (2004).

6. P. Fries, A mechanism for cognitive dynamics: neuronal communication through neuronal coherence. Trends Cogn Sci 9, 474–480 (2005).

7. T. Womelsdorf et al., Modulation of neuronal interactions through neuronal synchronization. Science 316, 1609–1612 (2007).

8. G. Laurent, Olfactory network dynamics and the coding of multidimensional signals. Nat Rev Neurosci 3, 884–895 (2002).

9. G. G. Gregoriou, S. J. Gotts, H. Zhou, R. Desimone, High-frequency, long-range coupling between prefrontal and visual cortex during attention. Science 324, 1207–1210 (2009).

10. T. Akam, D. M. Kullmann, Oscillations and filtering networks support flexible routing of information. Neuron 67, 308–320 (2010).

11. C. A. Bosman et al., Attentional stimulus selection through selective synchronization between monkey visual areas. Neuron 75, 875–888 (2012).

12. P. Fries, Rhythms for Cognition: Communication through Coherence. Neuron 88, 220–235 (2015).

13. G. Hahn, A. F. Bujan, Y. Fregnac, A. Aertsen, A. Kumar, Communication through resonance in spiking neuronal networks. PLoS Comput Biol 10, e1003811 (2014).

14. A. Palmigiano, T. Geisel, F. Wolf, D. Battaglia, Flexible information routing by transient synchrony. Nat Neurosci 20, 1014–1022 (2017).

15. L. L. Colgin et al., Frequency of gamma oscillations routes flow of information in the hippocampus. Nature 462, 353–357 (2009).

16. G. Buzsaki, Theta oscillations in the hippocampus. Neuron 33, 325–340 (2002).

17. J. O’Keefe, M. L. Recce, Phase relationship between hippocampal place units and the EEG theta rhythm. Hippocampus 3, 317–330 (1993).

18. M. E. Hasselmo, C. Bodelon, B. P. Wyble, A proposed function for hippocampal theta rhythm: separate phases of encoding and retrieval enhance reversal of prior learning. Neural Comput 14, 793–817 (2002).

19. J. Huxter, N. Burgess, J. O’Keefe, Independent rate and temporal coding in hippocampal pyramidal cells. Nature 425, 828–832 (2003).

20. L. L. Colgin, Mechanisms and functions of theta rhythms. Annu Rev Neurosci 36, 295–312 (2013).

21. G. Buzsaki, E. I. Moser, Memory, navigation and theta rhythm in the hippocampal-entorhinal system. Nat Neurosci 16, 130–138 (2013).

22. A. Z. Vollan, R. J. Gardner, M. B. Moser, E. I. Moser, Left-right-alternating theta sweeps in entorhinal-hippocampal maps of space. Nature 639, 995–1005 (2025).

23. A. Z. Vollan, M. F. Schellenberger, R. J. Gardner, M. B. Moser, E. I. Moser, Adaptive modulation of theta sweeps in the brain’s navigation circuit. Science 393, eaef4184 (2026).

24. M. Wang, D. J. Foster, B. E. Pfeiffer, Alternating sequences of future and past behavior encoded within hippocampal theta oscillations. Science 370, 247–250 (2020).

25. M. W. Jones, M. A. Wilson, Theta rhythms coordinate hippocampal-prefrontal interactions in a spatial memory task. PLoS Biol 3, e402 (2005).

26. A. G. Siapas, E. V. Lubenov, M. A. Wilson, Prefrontal phase locking to hippocampal theta oscillations. Neuron 46, 141–151 (2005).

27. J. M. Hyman, E. A. Zilli, A. M. Paley, M. E. Hasselmo, Working Memory Performance Correlates with Prefrontal-Hippocampal Theta Interactions but not with Prefrontal Neuron Firing Rates. Front Integr Neurosci 4, 2 (2010).

28. K. Benchenane et al., Coherent theta oscillations and reorganization of spike timing in the hippocampal-prefrontal network upon learning. Neuron 66, 921–936 (2010).

29. H. L. Hallock, A. Wang, A. L. Griffin, Ventral Midline Thalamus Is Critical for Hippocampal-Prefrontal Synchrony and Spatial Working Memory. J Neurosci 36, 8372–8389 (2016).

30. H. T. Ito, E. I. Moser, M. B. Moser, Supramammillary Nucleus Modulates Spike-Time Coordination in the Prefrontal-Thalamo-Hippocampal Circuit during Navigation. Neuron 99, 576–587 e575 (2018).

31. H. T. Ito, S. J. Zhang, M. P. Witter, E. I. Moser, M. B. Moser, A prefrontal-thalamo-hippocampal circuit for goal-directed spatial navigation. Nature 522, 50–55 (2015).

32. A. B. Tort et al., Dynamic cross-frequency couplings of local field potential oscillations in rat striatum and hippocampus during performance of a T-maze task. Proc Natl Acad Sci U S A 105, 20517–20522 (2008).

33. M. A. van der Meer, A. D. Redish, Theta phase precession in rat ventral striatum links place and reward information. J Neurosci 31, 2843–2854 (2011).

34. R. P. Vertes, B. Kocsis, Brainstem-diencephalo-septohippocampal systems controlling the theta rhythm of the hippocampus. Neuroscience 81, 893–926 (1997).

35. J. N. Rawlins, J. Feldon, J. A. Gray, Septo-hippocampal connections and the hippocampal theta rhythm. Exp Brain Res 37, 49–63 (1979).

36. I. Zutshi et al., Hippocampal Neural Circuits Respond to Optogenetic Pacing of Theta Frequencies by Generating Accelerated Oscillation Frequencies. Current biology : CB 28, 1179–1188 e1173 (2018).

37. B. Kocsis, R. P. Vertes, Characterization of neurons of the supramammillary nucleus and mammillary body that discharge rhythmically with the hippocampal theta rhythm in the rat. J Neurosci 14, 7040–7052 (1994).

38. B. Kocsis, R. P. Vertes, Phase relations of rhythmic neuronal firing in the supramammillary nucleus and mammillary body to the hippocampal theta activity in urethane anesthetized rats. Hippocampus 7, 204–214 (1997).

39. W. X. Pan, N. McNaughton, The medial supramammillary nucleus, spatial learning and the frequency of hippocampal theta activity. Brain Res 764, 101–108 (1997).

40. I. J. Kirk, Frequency modulation of hippocampal theta by the supramammillary nucleus, and other hypothalamo-hippocampal interactions: mechanisms and functional implications. Neurosci Biobehav Rev 22, 291–302 (1998).

41. W. X. Pan, N. McNaughton, The supramammillary area: its organization, functions and relationship to the hippocampus. Progress in neurobiology 74, 127–166 (2004).

42. J. S. Thinschmidt, G. G. Kinney, B. Kocsis, The supramammillary nucleus: is it necessary for the mediation of hippocampal theta rhythm? Neuroscience 67, 301–312 (1995).

43. W. X. Pan, N. McNaughton, The role of the medial supramammillary nucleus in the control of hippocampal theta activity and behaviour in rats. Eur J Neurosci 16, 1797–1809 (2002).

44. R. Kramis, C. H. Vanderwolf, B. H. Bland, Two types of hippocampal rhythmical slow activity in both the rabbit and the rat: relations to behavior and effects of atropine, diethyl ether, urethane, and pentobarbital. Exp Neurol 49, 58–85 (1975).

45. B. Schmidt et al., Dissociation between dorsal and ventral hippocampal theta oscillations during decision-making. J Neurosci 33, 6212–6224 (2013).

46. E. Kropff, J. E. Carmichael, E. I. Moser, M. B. Moser, Frequency of theta rhythm is controlled by acceleration, but not speed, in running rats. Neuron 109, 1029–1039 e1028 (2021).

47. J. P. Kennedy et al., A Direct Comparison of Theta Power and Frequency to Speed and Acceleration. J Neurosci 42, 4326–4341 (2022).

48. I. Daubechies, J. Lu, H.-T. Wu, Synchrosqueezed Wavelet Transforms: a Tool for Empirical Mode Decomposition. Appl. Comput. Harmon. Anal. 30, 243–261 (2011).

49. J. R. Climer, R. DiTullio, E. L. Newman, M. E. Hasselmo, U. T. Eden, Examination of rhythmicity of extracellularly recorded neurons in the entorhinal cortex. Hippocampus 25, 460–473 (2015).

50. U. Rutishauser, I. B. Ross, A. N. Mamelak, E. M. Schuman, Human memory strength is predicted by theta-frequency phase-locking of single neurons. Nature 464, 903–907 (2010).

51. P. Fries, J. H. Reynolds, A. E. Rorie, R. Desimone, Modulation of oscillatory neuronal synchronization by selective visual attention. Science 291, 1560–1563 (2001).

52. T. Hayakawa, H. Ito, K. Zyo, Neuroanatomical study of afferent projections to the supramammillary nucleus of the rat. Anat Embryol (Berl) 188, 139–148 (1993).

53. M. Mahn et al., Efficient optogenetic silencing of neurotransmitter release with a mosquito rhodopsin. Neuron 109, 1621–1635 e1628 (2021).

54. R. P. Vertes, J. T. McKenna, Collateral projections from the supramammillary nucleus to the medial septum and hippocampus. Synapse 38, 281–293 (2000).

55. Y. Hashimotodani, F. Karube, Y. Yanagawa, F. Fujiyama, M. Kano, Supramammillary Nucleus Afferents to the Dentate Gyrus Co-release Glutamate and GABA and Potentiate Granule Cell Output. Cell Rep 25, 2704–2715 e2704 (2018).

56. C. Nyakas, P. G. Luiten, D. G. Spencer, J. Traber, Detailed projection patterns of septal and diagonal band efferents to the hippocampus in the rat with emphasis on innervation of CA1 and dentate gyrus. Brain Res Bull 18, 533–545 (1987).

57. R. P. Gaykema, P. G. Luiten, C. Nyakas, J. Traber, Cortical projection patterns of the medial septum-diagonal band complex. J Comp Neurol 293, 103–124 (1990).

58. J. A. Ainge, M. A. van der Meer, R. F. Langston, E. R. Wood, Exploring the role of context-dependent hippocampal activity in spatial alternation behavior. Hippocampus 17, 988–1002 (2007).

59. A. Treisman, The binding problem. Curr Opin Neurobiol 6, 171–178 (1996).

60. C. von der Malsburg, W. Schneider, A neural cocktail-party processor. Biol Cybern 54, 29–40 (1986).

61. S. Ray, J. H. Maunsell, Differences in gamma frequencies across visual cortex restrict their possible use in computation. Neuron 67, 885–896 (2010).

62. M. Schneider et al., A mechanism for inter-areal coherence through communication based on connectivity and oscillatory power. Neuron 109, 4050–4067 e4012 (2021).

63. S. Ray, J. H. Maunsell, Do gamma oscillations play a role in cerebral cortex? Trends Cogn Sci 19, 78–85 (2015).

64. P. T. Huerta, J. E. Lisman, Heightened synaptic plasticity of hippocampal CA1 neurons during a cholinergically induced rhythmic state. Nature 364, 723–725 (1993).

65. I. J. Kirk, N. McNaughton, Mapping the differential effects of procaine on frequency and amplitude of reticularly elicited hippocampal rhythmical slow activity. Hippocampus 3, 517–525 (1993).

66. S. J. Mitchell, J. N. Rawlins, O. Steward, D. S. Olton, Medial septal area lesions disrupt theta rhythm and cholinergic staining in medial entorhinal cortex and produce impaired radial arm maze behavior in rats. J Neurosci 2, 292–302 (1982).

67. N. McNaughton et al., Contribution of synapses in the medial supramammillary nucleus to the frequency of hippocampal theta rhythm in freely moving rats. Hippocampus 5, 534–545 (1995).

68. J. S. Farrell et al., Supramammillary regulation of locomotion and hippocampal activity. Science 374, 1492–1496 (2021).

69. J. Kiss, A. Csaki, H. Bokor, K. Kocsis, G. Szeiffert, Topographic localization of calretinin, calbindin, VIP, substance P, CCK and metabotropic glutamate receptor immunoreactive neurons in the supramammillary and related areas of the rat. Neurobiology (Bp) 5, 361–388 (1997).

70. J. M. Burgunder, W. S. Young, 3rd, Neurons containing cholecystokinin mRNA in the mammillary region: ontogeny and adult distribution in the rat. Cell Mol Neurobiol 9, 281–294 (1989).

71. E. R. Wood, P. A. Dudchenko, R. J. Robitsek, H. Eichenbaum, Hippocampal neurons encode information about different types of memory episodes occurring in the same location. Neuron 27, 623–633 (2000).

72. L. M. Frank, E. N. Brown, M. Wilson, Trajectory encoding in the hippocampus and entorhinal cortex. Neuron 27, 169–178 (2000).

73. R. P. Vertes, PHA-L analysis of projections from the supramammillary nucleus in the rat. J Comp Neurol 326, 595–622 (1992).

74. J. J. Jun et al., Fully integrated silicon probes for high-density recording of neural activity. Nature 551, 232–236 (2017).

75. N. A. Steinmetz et al., Neuropixels 2.0: A miniaturized high-density probe for stable, long-term brain recordings. Science 372, (2021).

76. M. Pachitariu, S. Sridhar, J. Pennington, C. Stringer, Spike sorting with Kilosort4. Nat Methods 21, 914–921 (2024).

